# Formal model of Parkinson’s disease neurons unveils possible causality links in the pathophysiology of the disease

**DOI:** 10.1101/2020.02.06.937664

**Authors:** Morgane Nadal, Gabriele S. Kaminski Schierle, Duygu Dikicioglu

**Affiliations:** Department of Biology, Ecole Normale Supérieure – PSL University, 75005, Paris, France; Department of Biotechnology and Chemical Engineering, University of Cambridge, CB3 0AS, Cambridge, United Kingdom; Department of Biochemical Engineering, University College London, WC1E 6BT, London, UK

**Keywords:** Parkinson, neurodegenerative disease, systems biology, causality, dopaminergic neuron, α-synuclein, iron homeostasis, ROS, petri nets

## Abstract

Parkinson’s Disease is the second most common neurodegenerative disease after Alzheimer’s disease. Despite extensive research, the initial cause of the disease is still unknown, although substantial advances were made in understanding of its genetics and the cognate neurophysiological mechanisms. Determining the causality relationships and the chronological steps pertaining to Parkinson’s Disease is essential for the discovery of novel drug targets. We developed a systematic *in silico* model based on available data, which puts the possible sequence of events occurring in a neuron during disease onset into light. This is the first ever attempt, to our knowledge, to model comprehensively the primary modifications in the molecular pathways that manifest in compromised neurons from the commencement of the disease to the consequences of its progression. We showed that our proposed disease pathway was relevant for unveiling yet incomplete knowledge on calcium homeostasis in mitochondria, ROS production and α-synuclein misfolding.

**Graphical abstract:** 

**Highlights:**

- Varying calcium concentration in aging dopaminergic neurons triggers disease onset.
- ROS production in the mitochondria potentially causes iron accumulation.
- Iron homeostasis dysregulation is linked to α-synuclein aggregation.

## Introduction

Parkinson’s Disease (PD) affects about 6.3 million people worldwide (EBC, 2015), with a prevalence of about 3% for the population aged over 80 years. PD is characterized by bradykinesia, rigidity and tremor, but can be accompanied by non-motor symptoms such as cognitive impairment and depression (Poewe et al., 2017). The initial cause of the disease is still unknown, but considerable advances have been made in the understanding of genetics and neurophysiological mechanisms underlying the disease and its symptoms. The pathophysiology of PD has recently been put forward by the discovery of many genes involved in rare forms of the disease, which brings the main pathways associated with PD to light (Fujita et al., 2014; Singleton et al., 2013; Tysnes and Storstein, 2017). The characteristic features of PD are localized loss of Substantia Nigra pars compacta dopaminergic neurons (SNpc DA neurons), mitochondrial dysfunction accompanied by oxidative stress, accumulation of free iron (Mochizuki and Yasuda, 2012; Salvador, 2010) and accumulation and prion-like propagation of the misfolded α-synuclein protein (Poewe et al., 2017).

The autonomous pacemaking activity of SNpc DA neurons is linked to cyclic variations in calcium concentration in a range of 0.1 - 2-3 µM, and up to >10 µM in microdomains within the cell (Raffaello et al., 2016; Surmeier et al., 2010). In young individuals, pacemaking relies on Na^+^ channels, which are activated by hyperpolarization. With age, this activity is increasingly driven by voltage-dependent L-type calcium channels, composed of a specific voltage-gated pore-forming subunit (Cav1.3) (Sulzer and Surmeier, 2013; Surmeier and Schumacker, 2013). Blocking these channels were shown to reduce the risk of PD (Chan et al., 2007). The energy cost of pacemaking is high; more than 50% and up to 75% of the total ATP produced was reported to be required for maintaining the ions gradients at the cellular level (Duda et al., 2016; Mamelak, 2018). Calcium interacts with multiple cell components and can be stored in buffers or organelles such as the ER and the mitochondria. Calcium accelerates the energy metabolism, usually by binding to the Krebs cycle enzymes, upon entering mitochondria (Tarasov et al., 2012). This acceleration can lead to the accumulation of reactive oxygen species (ROS) in the mitochondria and possibly cause oxidative stress. Indeed, mitochondria are thought to work at maximal capacity and have limited energy reserves, facilitating the manifestation of any potential dysfunction accompanied by its rapid amplification in the neuron (Mamelak, 2018). In addition, the reduction of the activity of complex I of the electron transfer chain (ETC) has also been observed in early stages in disease-advanced patients, as well as being a main cause of early parkinsonism (Langston et al., 1983). ROS production was reported to be aggravated when the ETC was blocked at key sites, including those in complex I (Dias et al., 2013; Zhao et al., 2019).

Iron was shown to accumulate in PD neurons twice as much as that in healthy brains (Sian□Hülsmann et al., 2011), however the mechanisms of its accumulation are still unknown. At late stages of the disease, almost all patients are subject to an intraneuronal aggregation of the α-synuclein protein (Halliday et al., 2011). α-synuclein was shown to aggregate inside the cytosol due to its overabundance and misfolding. These proteins were shown to propagate in the brain in a prion-like fashion (Poewe et al., 2017). The function of α-synuclein, a monomeric 140 amino-acid protein encoded by the SNCA gene, is not fully understood but the protein appears to be essential for the neuron. Multiple interactions were identified or were reported likely to exist between this intrinsically disordered soluble protein and cellular, mitochondrial, lysosomal and vesicular membranes, Ca^2+^ and Fe^2+^ cations, enzymes, proteins, and the cytoskeleton among others (Faustini et al., 2017; Lautenschläger et al., 2017; Poewe et al., 2017; Post et al., 2018; Stephens et al., 2018). The recent discovery of an iron response element on α-synuclein mRNA (Lingor et al., 2017; Rogers et al., 2011) implied that the synthesis of α-synuclein could potentially be triggered by a variation in cytosolic iron concentration.

In order to employ a systematic approach to decipher the complexity of the molecular pathways involved in PD neurons, we constructed an *in silico* model of a neuron developing Parkinson’s disease. We constructed and tested the model, based on biological data available in the literature. Our first proposition was that the autonomous pacemaking led the cell to an unstable equilibrium where a small dysfunction would enforce neuronal degeneration, due to the high rate of metabolic activity and the limited availability of energy reserves. Second, an age-associated dysfunctional autonomous pacemaking triggered an increase in ROS production. Third, ROS concentration damaged cellular functionality, particularly disrupted iron homeostasis leading to the accumulation of free iron. Finally, we propose that the increase in free iron concentration would be sufficient to trigger the translation of α-synuclein at a high rate. All these propositions were shown to be revenant for utilizing this model to unveil knowledge gaps in mitochondrial calcium homeostasis, ROS production and α-synuclein misfolding pertaining to the onset and progression of the disease.

## Results

We structured our model around four successive biological “modules” each representing a process related to Parkinson’s disease: (i) calcium metabolism, (ii) energy and oxidative metabolism, (iii) iron homeostasis, and (iv) α-synuclein misfolding and aggregation (Figure 1.A.). Each module was first constructed and tested separately using the available data. Petri Net formalism was employed in constructing and simulating the model substructures. We employed number of molecules of each entity at the cellular level or in different cellular compartments in order to facilitate its construction. Regulatory mechanisms were proposed and included in the model as necessary in order to compensate for missing knowledge and data unavailability. We discuss each module below in light of these aspects.

**Figure 1:**
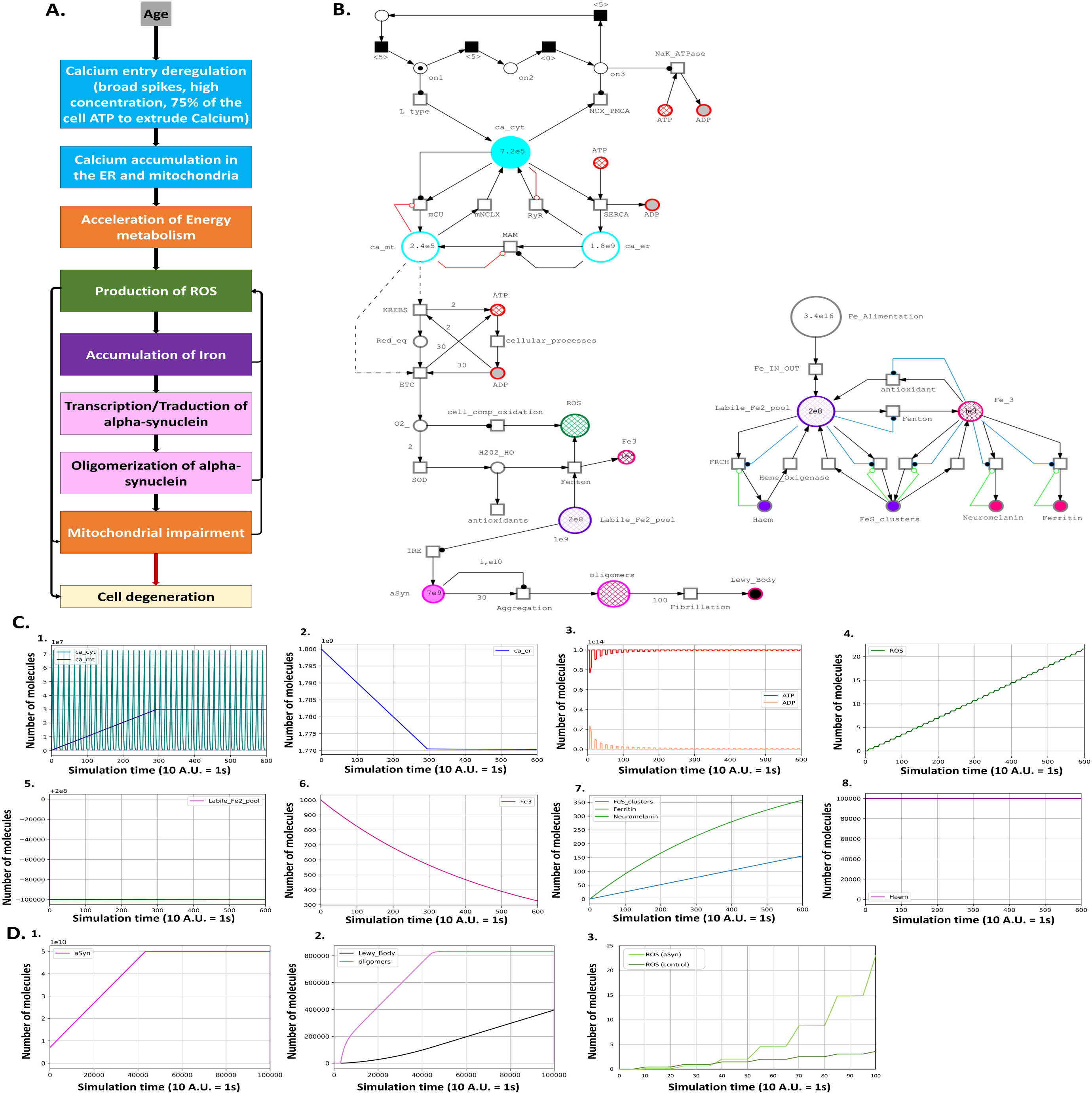
A. Simplified Modules representing a possible and plausible time sequence of events until SNpc DA neuron death in PD. Each module represents a pathway implicated in the disease. The arrows represent the relationships and links between the different cell mechanisms. For details, refer to Figure 2 and the Results section for the construction steps of the modules. **B. SNpc DA neuron Petri Net.** The five previously defined modules; calcium metabolism, energy metabolism, ROS production, iron metabolism and α-synuclein aggregation compiled together. **C. Evolution of the molecular species number in a SNpc DA neuron during the first 1 min of life.** 1 A.U. = 1 sec. **1.** Cytoplasmic and mitochondrial calcium. **2.** ER calcium. **3.** ATP and ADP. **4**. ROS. **5.** Fe^2+^. **6.** Fe^3+^. **7.** Ferritin, Neuromelanin and Fe–S clusters. Neuromelanin and ferritin curves are overlapping. **8.** Haem. For more details refer to the main text. **D. Case Studies. 1.2. Evolution of the quantity of α-synuclein, oligomers and Lewy Bodies.** 1 A.U. = 1 sec. **1.** α-synuclein is translated linearly from 7.10^9^ molecules (control and initial concentration) to 5.10^10^ molecules. **2.** Once α-synuclein has reached the 1.10^10^ molecules threshold, oligomers start to form and the number of oligomers reaches about 8.10^5^ molecules after 2 days. This is followed by the formation of Lewy Bodies/fibrils, which reaches 4.10^5^ entities. **3.** α**-synuclein effect on ROS accumulation.** The accumulation of ROS without the intervention of α-synuclein is globally linear and reaches 4 molecules in 10 sec. The accumulation of ROS with the action of α-synuclein is exponential and reaches 24 molecules in 10 sec.

### Calcium-mediated autonomous pacemaking

We considered this module to comprise of two tasks, namely calcium uptake and release at the membrane, and calcium buffering in the cell.

We grouped the key channel elements in modelling membrane-associated events. The T- and L-type channels for calcium uptake were modelled together by approximating the calcium entry rate as the difference between the initial and final calcium concentration in the cell over the duration of the channel opening (Branch et al., 2014; Duda et al., 2016; Poetschke et al., 2015). For calcium release, we considered NCX and PMCA pumps and modeled the exit of calcium based on mass action kinetics in order to achieve the empirically expected spiked calcium concentration profile. Simulation of the model yielded the cyclic variations of calcium in the cytosol, correlating with the neuron pacemaking activity with spikes of 1 ms (10 U.A., Figure 1.B.). This model of pacemaking was then adapted to an ageing individual. It has been shown that aged individuals present more stochasticity in the autonomous pacemaking in their SNpc DA neurons manifesting as reduction in the control of delay between the spikes (Branch et al., 2014, 2016), which we modelled utilising stochastic transitions in the uptake and release of calcium (for details see STAR methods).

Calcium storage was only considered in the ER and in the mitochondria for modelling the buffering of calcium in the neuron (for details regarding the channels, see Figure 2 and Table 1 in STAR Methods) because of the prevalence of this activity in these subcellular compartments (Raffaello et al., 2016). A threshold concentration for calcium transport into the mitochondria was selected arbitrarily as 10 μM since this is the extramitochondrial concentration that allows the channels to open and let calcium flow in (Rizzuto and Pozzan, 2006; Surmeier et al., 2010; Zaichick et al., 2017). This artificial upper bound was ensured through an inhibitory arc (Figure 1.B. and Figure S1.a.) in order to avoid unbounded increase of mitochondrial calcium concentration, which is not biologically relevant. This artificial intervention possibly denoted a yet unreported regulatory mechanism of feedback inhibition; its empirical investigation would indeed improve our understanding of calcium homeostasis. Simulation of the model indicates that a steady state is rapidly attained for the concentration of calcium in the ER and the mitochondria. Our current understanding cannot reveal whether this is indeed the empirically observed phenomenon or whether oscillations are present, this remains yet an open question for empirical analysis proposed by our model.

**STAR Table 1:**
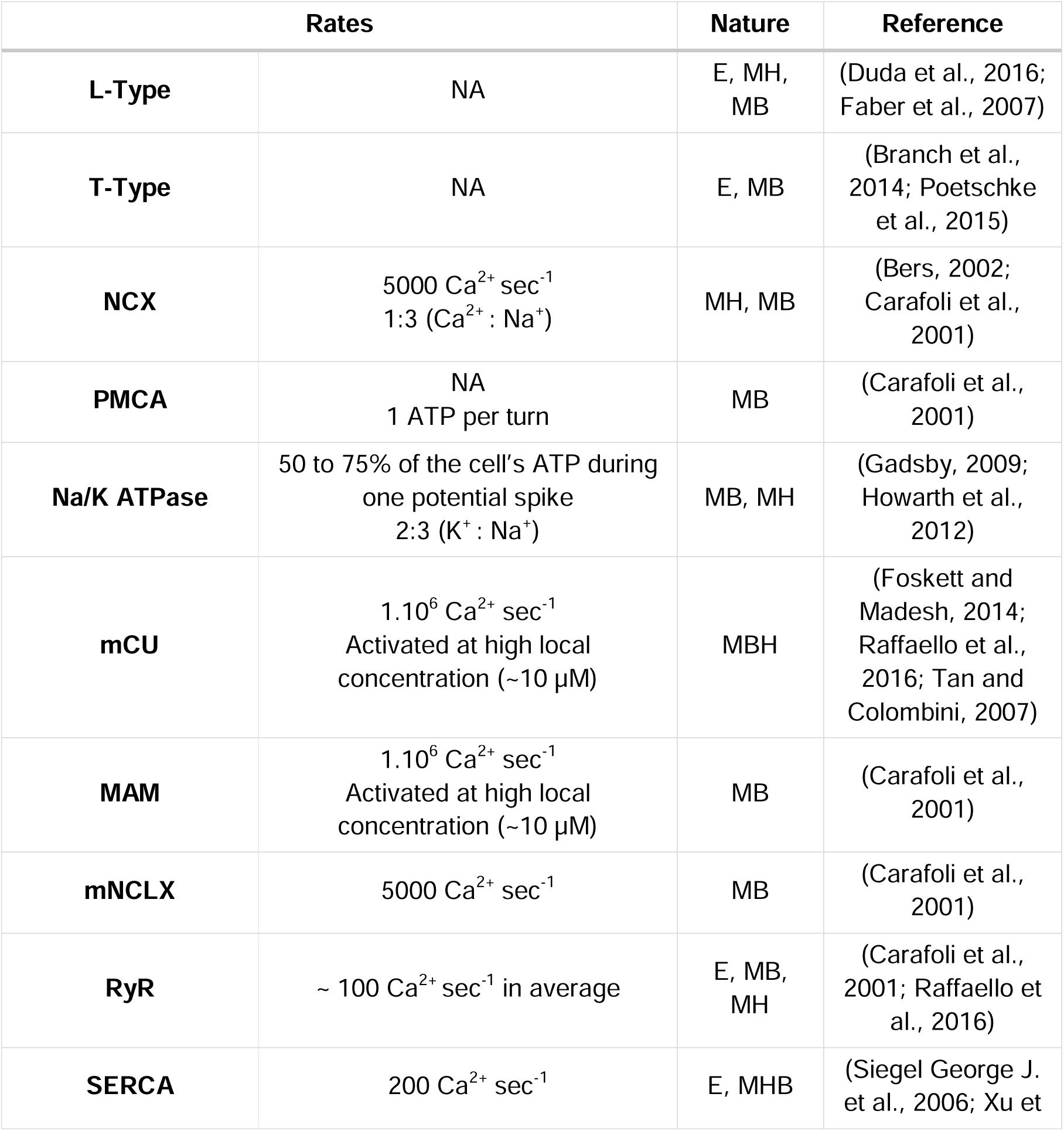
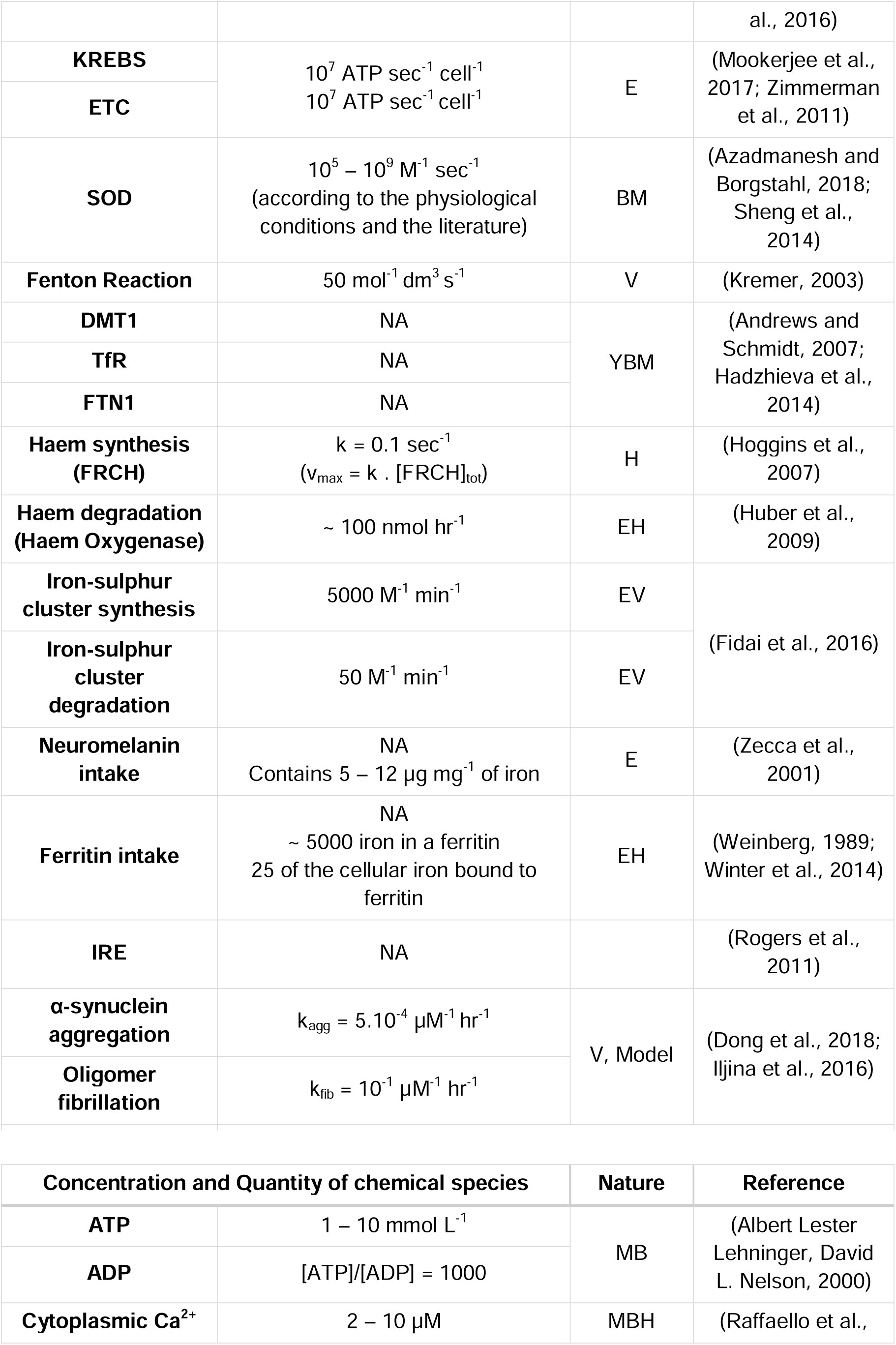
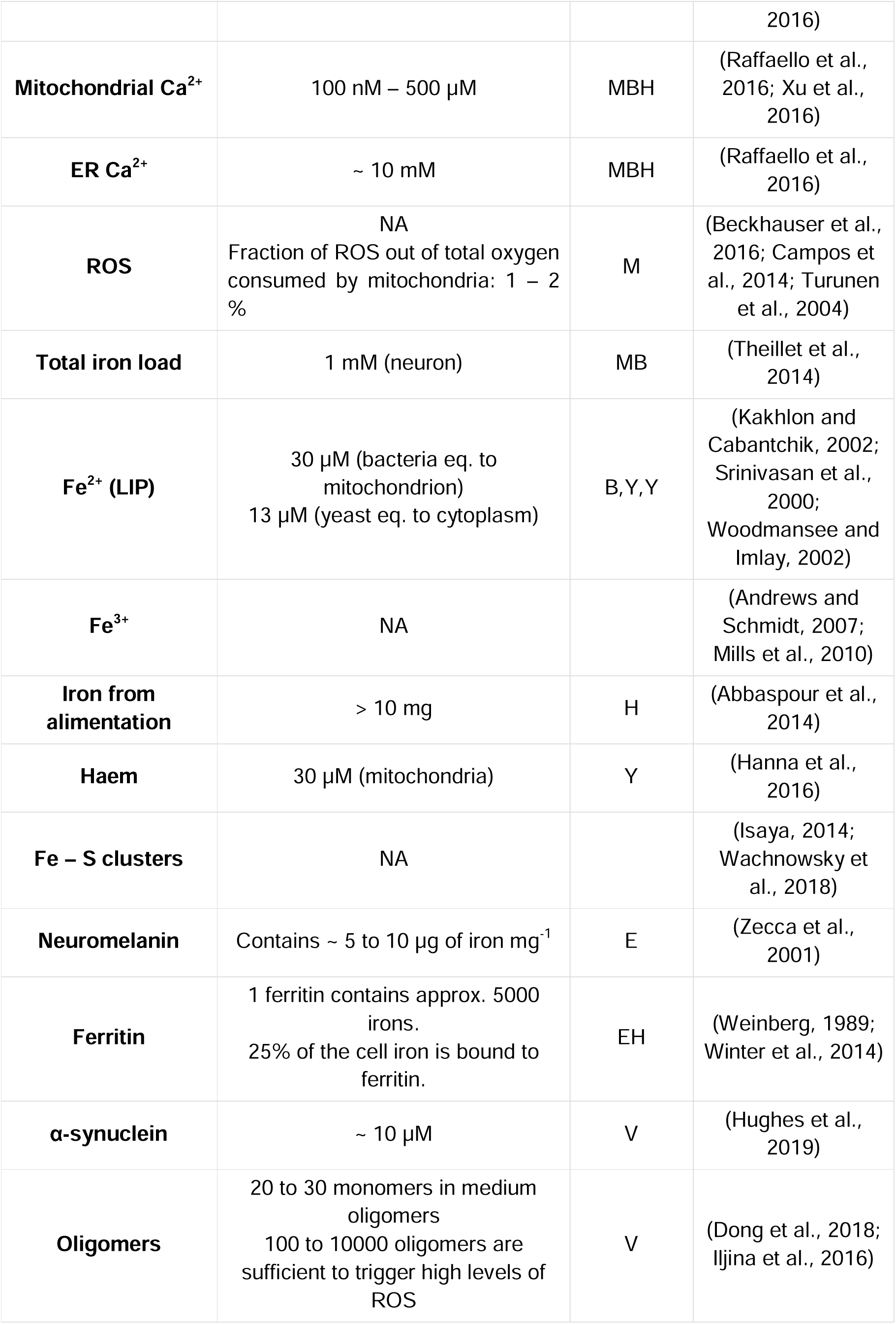
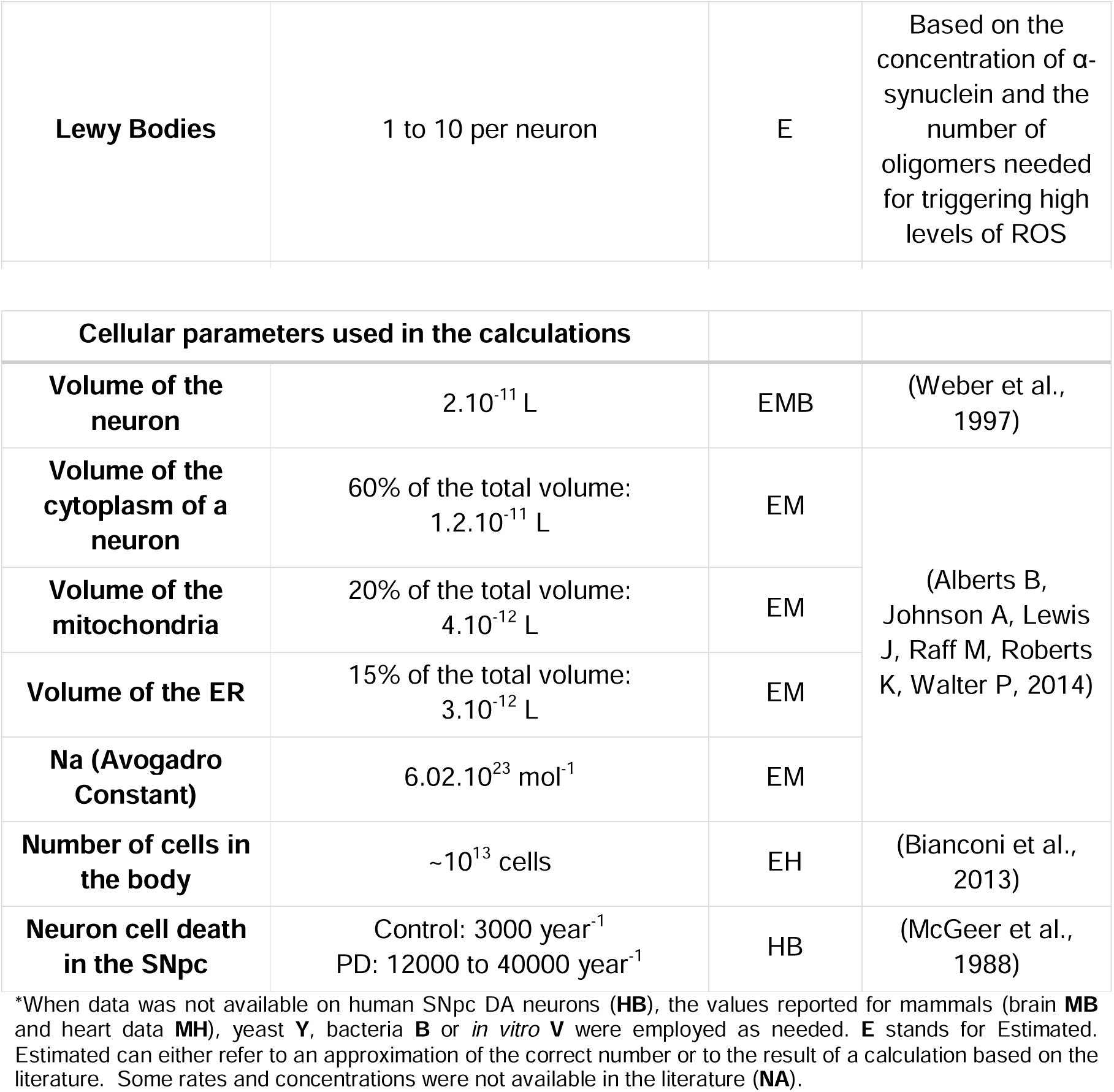
Rates of transitions and the number of molecules of model components*.

|  | **Rates** | **Nature** | **Reference** |
| --- | --- | --- | --- |
| **L-Type** | NA | E, MH, MB | (Duda et al., 2016; Faber et al., 2007) |
| **T-Type** | NA | E, MB | (Branch et al., 2014; Poetschke et al., 2015) |
| **NCX** | 5000 $\text{Ca}^{2+} \text{ sec}^{-1}$<br>1:3 ($\text{Ca}^{2+} : \text{Na}^{+}$) | MH, MB | (Bers, 2002; Carafoli et al., 2001) |
| **PMCA** | NA<br>1 ATP per turn | MB | (Carafoli et al., 2001) |
| **Na/K ATPase** | 50 to 75% of the cell's ATP during one potential spike<br>2:3 ($\text{K}^{+} : \text{Na}^{+}$) | MB, MH | (Gadsby, 2009; Howarth et al., 2012) |
| **mCU** | $1.10^6 \text{ Ca}^{2+} \text{ sec}^{-1}$<br>Activated at high local concentration ($\sim 10 \mu\text{M}$) | MBH | (Foskett and Madesh, 2014; Raffaello et al., 2016; Tan and Colombini, 2007) |
| **MAM** | $1.10^6 \text{ Ca}^{2+} \text{ sec}^{-1}$<br>Activated at high local concentration ($\sim 10 \mu\text{M}$) | MB | (Carafoli et al., 2001) |
| **mNCLX** | 5000 $\text{Ca}^{2+} \text{ sec}^{-1}$ | MB | (Carafoli et al., 2001) |
| **RyR** | $\sim 100 \text{ Ca}^{2+} \text{ sec}^{-1}$ in average | E, MB, MH | (Carafoli et al., 2001; Raffaello et al., 2016) |
| **SERCA** | 200 $\text{Ca}^{2+} \text{ sec}^{-1}$ | E, MHB | (Siegel George J. et al., 2006; Xu et al., 2016) |
| **KREBS** | $10^7 \text{ ATP sec}^{-1} \text{ cell}^{-1}$<br>$10^7 \text{ ATP sec}^{-1} \text{ cell}^{-1}$ | E | (Mookerjee et al., 2017; Zimmerman et al., 2011) |
| **ETC** |  |  |  |
| **SOD** | $10^5 - 10^9 \text{ M}^{-1} \text{ sec}^{-1}$<br>(according to the physiological conditions and the literature) | BM | (Azadmanesh and Borgstahl, 2018; Sheng et al., 2014) |
| **Fenton Reaction** | $50 \text{ mol}^{-1} \text{ dm}^3 \text{ s}^{-1}$ | V | (Kremer, 2003) |
| **DMT1** | NA | YBM | (Andrews and Schmidt, 2007; Hadzhieva et al., 2014) |
| **TfR** | NA |  |  |
| **FTN1** | NA |  |  |
| **Haem synthesis (FRCH)** | $k = 0.1 \text{ sec}^{-1}$<br>($v_{\max} = k \cdot [\text{FRCH}]_{\text{tot}}$) | H | (Hoggins et al., 2007) |
| **Haem degradation (Haem Oxygenase)** | $\sim 100 \text{ nmol hr}^{-1}$ | EH | (Huber et al., 2009) |
| **Iron-sulphur cluster synthesis** | $5000 \text{ M}^{-1} \text{ min}^{-1}$ | EV | (Fidai et al., 2016) |
| **Iron-sulphur cluster degradation** | $50 \text{ M}^{-1} \text{ min}^{-1}$ | EV | |
| **Neuromelanin intake** | NA<br>Contains $5 - 12 \mu\text{g mg}^{-1}$ of iron | E | (Zecca et al., 2001) |
| **Ferritin intake** | NA<br>$\sim 5000$ iron in a ferritin<br>25 of the cellular iron bound to ferritin | EH | (Weinberg, 1989; Winter et al., 2014) |
| **IRE** | NA |  | (Rogers et al., 2011) |
| **α-synuclein aggregation** | $k_{\text{agg}} = 5 \cdot 10^{-4} \mu\text{M}^{-1} \text{ hr}^{-1}$ | V, Model | (Dong et al., 2018; Iljina et al., 2016) |
| **Oligomer fibrillation** | $k_{\text{fib}} = 10^{-1} \mu\text{M}^{-1} \text{ hr}^{-1}$ | | |

| Concentration and Quantity of chemical species |  | Nature | Reference |
| --- | --- | --- | --- |
| **ATP** | $1 - 10 \text{ mmol L}^{-1}$ | MB | (Albert Lester Lehninger, David L. Nelson, 2000) |
| **ADP** | $[\text{ATP}]/[\text{ADP}] = 1000$ | | |
| **Cytoplasmic $\text{Ca}^{2+}$** | $2 - 10 \mu\text{M}$ | MBH | (Raffaello et al., |
|  |  |  | 2016) |
| **Mitochondrial Ca<sup>2+</sup>** | 100 nM – 500 µM | MBH | (Raffaello et al., 2016; Xu et al., 2016) |
| **ER Ca<sup>2+</sup>** | ~ 10 mM | MBH | (Raffaello et al., 2016) |
| **ROS** | NA<br>Fraction of ROS out of total oxygen consumed by mitochondria: 1 – 2 % | M | (Beckhauser et al., 2016; Campos et al., 2014; Turunen et al., 2004) |
| **Total iron load** | 1 mM (neuron) | MB | (Theillet et al., 2014) |
| **Fe<sup>2+</sup> (LIP)** | 30 µM (bacteria eq. to mitochondrion)<br>13 µM (yeast eq. to cytoplasm) | B,Y,Y | (Kakhlon and Cabantchik, 2002; Srinivasan et al., 2000; Woodmansee and Imlay, 2002) |
| **Fe<sup>3+</sup>** | NA |  | (Andrews and Schmidt, 2007; Mills et al., 2010) |
| **Iron from alimentation** | > 10 mg | H | (Abbaspour et al., 2014) |
| **Haem** | 30 µM (mitochondria) | Y | (Hanna et al., 2016) |
| **Fe – S clusters** | NA |  | (Isaya, 2014; Wachnowsky et al., 2018) |
| **Neuromelanin** | Contains ~ 5 to 10 µg of iron mg<sup>-1</sup> | E | (Zecca et al., 2001) |
| **Ferritin** | 1 ferritin contains approx. 5000 irons.<br>25% of the cell iron is bound to ferritin. | EH | (Weinberg, 1989; Winter et al., 2014) |
| **α-synuclein** | ~ 10 µM | V | (Hughes et al., 2019) |
| **Oligomers** | 20 to 30 monomers in medium oligomers<br>100 to 10000 oligomers are sufficient to trigger high levels of ROS | V | (Dong et al., 2018; Iljina et al., 2016) |
| **Lewy Bodies** | 1 to 10 per neuron | E | Based on the concentration of α-synuclein and the number of oligomers needed for triggering high levels of ROS |

| Cellular parameters used in the calculations |  |  |  |
| --- | --- | --- | --- |
| **Volume of the neuron** | $2 \cdot 10^{-11}$ L | EMB | (Weber et al., 1997) |
| **Volume of the cytoplasm of a neuron** | 60% of the total volume:<br>$1.2 \cdot 10^{-11}$ L | EM | (Alberts B, Johnson A, Lewis J, Raff M, Roberts K, Walter P, 2014) |
| **Volume of the mitochondria** | 20% of the total volume:<br>$4 \cdot 10^{-12}$ L | EM | |
| **Volume of the ER** | 15% of the total volume:<br>$3 \cdot 10^{-12}$ L | EM | |
| **Na (Avogadro Constant)** | $6.02 \cdot 10^{23}$ mol<sup>-1</sup> | EM | |
| **Number of cells in the body** | $\sim 10^{13}$ cells | EH | (Bianconi et al., 2013) |
| **Neuron cell death in the SNpc** | Control: 3000 year<sup>-1</sup><br>PD: 12000 to 40000 year<sup>-1</sup> | HB | (McGeer et al., 1988) |

\*When data was not available on human SNpc DA neurons (**HB**), the values reported for mammals (brain **MB** and heart data **MH**), yeast **Y**, bacteria **B** or *in vitro* **V** were employed as needed. **E** stands for Estimated. Estimated can either refer to an approximation of the correct number or to the result of a calculation based on the literature. Some rates and concentrations were not available in the literature (**NA**).

**Figure 2:**
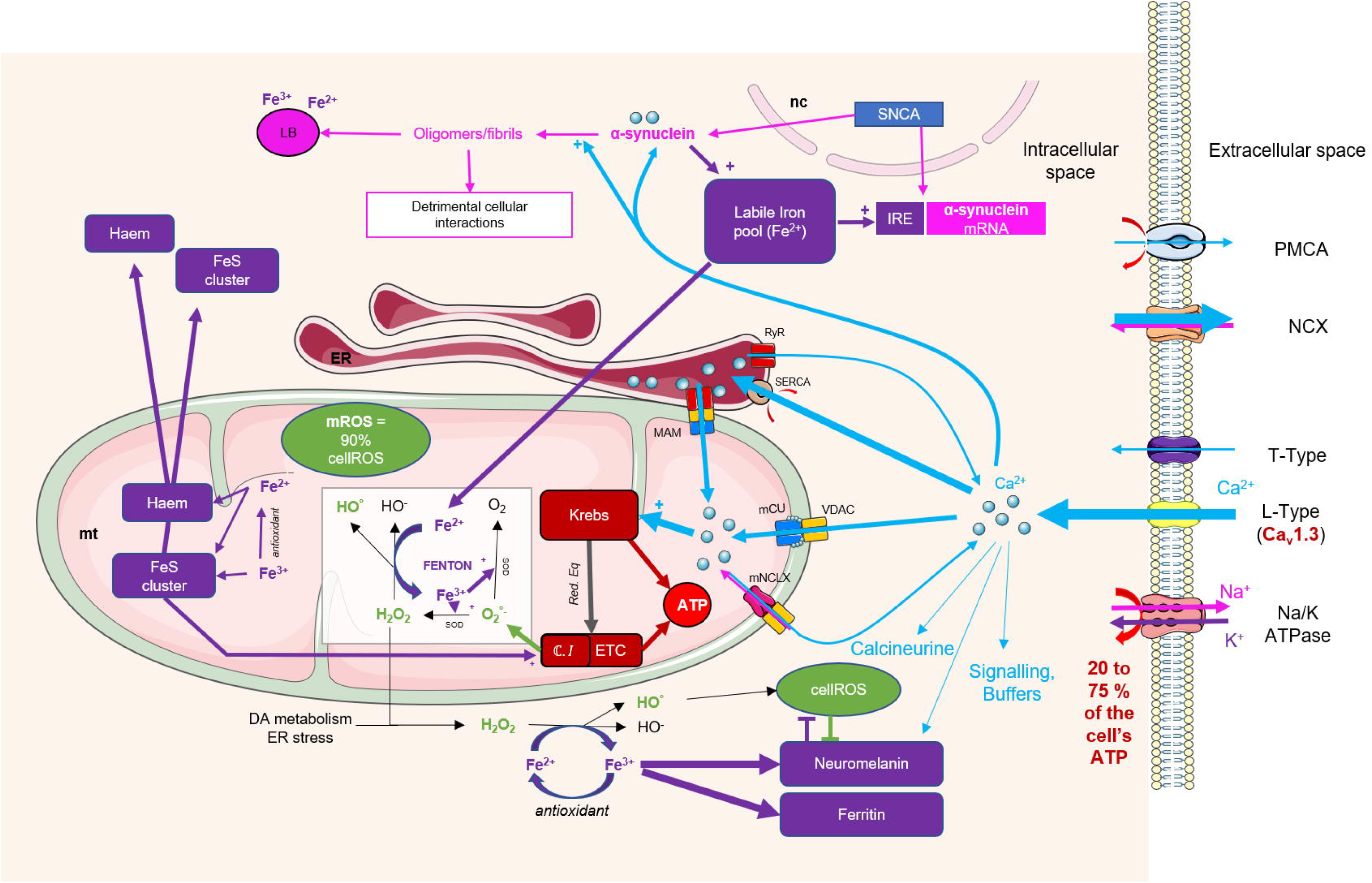
Schematic representation of the cellular system and its main mechanisms involved in PD. The autonomous pacemaking triggers cyclic entry of massive amounts of calcium into the SNpc DA neurons. The calcium can be used in signalling pathways, or be buffered in order to prevent from binding to detrimental cell components. Calcium is allowed to be transferred at a high rate to the ER and the mitochondria, and this transfer fosters the ATP synthesis by accelerating specific enzymes in the Krebs cycle. Acceleration of the energy metabolism renders more electrons susceptible to leak from the ETC. This generates ROS through the Fenton Reaction, with mitochondrial ROS being estimated to represent 90% of the total ROS produced by the cell. The Fenton Reaction implicates iron oxidation. Iron (III) can either be directly reduced by antioxidants or be stored in Ferritin or Neuromelanin. Iron (II) is essential for enzymatic activity and in the biosynthesis of Haems and FeS clusters. A large Labile Iron pool facilitates α-synuclein translation since the protein contains an IRE. An increase in α-synuclein concentration, which is normally highly controlled, may trigger its misfolding and aggregation in oligomers, fibrils and Lewy Bodies. The misfolded α-synuclein then interacts with cell components, and triggers, coupled with ROS, cell degeneration. Pathogenic α-synuclein can also propagate between cells as a prion. Arrows coupled with a plus represent activation. Flat-end arrows represent inhibition. mt: mitochondria. ER: Endoplasmic Reticulum. nc: nucleus. ETC: Electron Transfer Chain. SOD: Superoxide dismutase. ROS: Reactive Oxygen Species. IRE: Iron Response Element. LB: Lewy Body.

### Energy Metabolism and ROS production

The conservation of the number of ATP and ADP molecules and the electron leak from the ETC are aspects of energy generation, which needed to be considered in the model. The process was substantially simplified by considering the Krebs cycle as a single transition in order to avoid unnecessary complexity in the model. A similar approach was followed in modelling the ETC and its associated cellular processes. The critical parameters in this module were the rates of the proposed transitions representing clustered process steps. The rates were adjusted such that the conservation of the number of ATP and ADP molecules were ensured and these rates needed to be readjusted at every modification operated on the system, which involved a transaction of ATP. This challenge was overcome by setting an initial rate for the Krebs cycle and the ETC transitions and modify their prevalence in response to changing conditions through arc weights (for a detailed discussion on arc weights, see (Heiner et al., 2012; Rohr et al., 2010)). A constant number of ATP and ADP molecules could be maintained across time using this module, avoiding any oscillations or other perturbations in the consumption of ATP. It is important to note that this model considered the involvement of the Krebs cycle and the ETC in this specific process only, thus the nominal rates of transition employed in this model does not represent the energy requirements at the whole cell level. Nevertheless, it provides a reasonable and useful approximation for the segregation of process-specific energy requirements, which is not possible to determine empirically.

We then modelled two different paths of ROS production (Park et al., 2018; Trist et al., 2019) following superoxide (O_2_^°-^) synthesis. A fraction of the O_2_^°-^ can directly damage the cell components before being neutralised by cell defence systems. Alternatively, O2^°-^ is converted into hydrogen peroxide (H_2_O_2_) which can either dismute into unharmful radical species before it is neutralised by antioxidants or facilitates the iron-mediated Fenton Reaction. Mass action kinetics were employed throughout the module for simplicity since any data to contradict this assumption has not been reported elsewhere. This allowed linear ROS production as per a given amount of O_2_^°-^ molecules supplemented per arbitrary unit of time (A.U.). ROS production depended highly on the rate of neutralisation of the radicals and on the ETC activity, with high ROS rates accompanied by high ETC activity and low rates of neutralisation of the radicals by antioxidants.

### Iron homeostasis

Iron is implicated in the Fenton Reaction occurring in the mitochondria and in the cytoplasm, thus its homeostasis is tightly regulated. We had to simplify the molecular pathways involved in these mechanisms due to lack of available empirical data. We consider the total iron (II) and iron (III) available in the cell and don not distinguish between the mitochondrial, ER-associated or cytoplasmic iron (II). This simplification is valid for the purposes of this model since iron gets quickly transported in and out of the organelles without any energy requirements (Mills et al., 2010; Sian□Hülsmann et al., 2011). The labile iron (II) and iron (III) pools were ensured to possess non-negative values (blue arcs in Figure 1.B. and Figure S6). We also simplified the representation of haem biosynthesis as well as Fe-S cluster formation and maturation into a single transition each, also abstaining from the assignment of subcellular localisation of any intermediary steps of these processes. The Fenton Reaction and neutralisation of radicals by antioxidants represented all possible reactions leading to an equilibrium between iron (II), which is available at a much higher concentration in the cell, and iron (III), which is only found in minimal quantities. The channels and pumps (TfR, DMT, FTN) involved in the import and export of iron across the cellular boundary were represented by a single transition fuelled by the daily iron intake. Iron was always available for the system considering that daily intake would be sufficient to meet all cellular demands. The iron was taken up by the cell as iron (II), was then converted into iron (III) in the event of Fenton reaction taking place, or was incorporated into haem molecules and Fe–S clusters. Iron (III) was reduced to iron (II) by antioxidants or was stored in neuromelanin or ferritin (Mills et al., 2010; Núñez et al., 2014). We considered the accumulation of iron in neuromelanin as an irreversible but limited process. This assumption could be revisited in light of new discoveries related to the role of neuromelanin and ferritin in SNpc DA neurons in the future.

All iron species, with the exception of Fe-S clusters reached steady state within the first 500 seconds of the simulation. Ferritin and Neuromelanin production transitions were assigned the same rate and maximum cellular concentration available. The haem synthesis is almost instantaneous as indicated by the empirical rates, and all haem required for the model was synthesized within the first couple of seconds of the simulation. The number of Fe (III) molecules available in the neuronal cell at steady state was identified as very low (only 20 molecules), in line with early reports (Mackenzie et al., 2008; Ponka, 1999). The labile Fe (II) pool rapidly diminished within the initial seconds as Fe (II) was utilised in haem synthesis and this decrease slowed down as the haem concentration reached its steady state value.

### α-synuclein translation and aggregation

We propose a mechanism whereby iron accumulated during PD and triggered or activated the translation of α-synuclein when it reached a certain threshold, in light of the recent findings on the relationship between the IRE on the protein, and consequently model the onset of α-synuclein misfolding, aggregation and fibrillation accordingly. Moreover, α-synuclein was reported to accelerate the accumulation of iron, further aggravating the problem as proposed by our model (Davies et al., 2011; Lingor et al., 2017).

In this module, we will first consider the transition that represents the translation of α-synuclein. The concentration of α-synuclein is approximately 10 μM in the neurons of healthy individuals (Cremades et al., 2012). We propose that it is a tightly regulated mechanism and that a slight increase in α-synuclein concentration would initiate oligomerization in the model. We propose an arbitrary rate of α-synuclein translation since no empirical rate has been reported in the literature. An oligomer constitutes of approximately 30 α-synuclein molecules (Dong et al., 2018), and we proposed that 100 oligomers can turn a fibril into a Lewy Body. These reactions were modelled to be irreversible. We particularly focused on oligomers representing pathogenic elements in our model; they interacted with various cell components. For example, α-synuclein oligomers can accelerate ROS production; it was reported that 100 to 10000 oligomers were sufficient to generate high levels of ROS in PD neurons (Dong et al., 2018; Iljina et al., 2016).

The number of oligomers and Lewy Bodies increased proportionally as the number of α-synuclein proteins increase. Although we employed an arbitrary translation rate to better visualize the oligomerisation dynamics, any empirically determined rate sufficiently high to allow the initiation of α-synuclein translation would cause the net to display similar behaviour, albeit with a different time delay.

### Assembling the modules together: From autonomous pacemaking in the neuron to α-synuclein aggregation

Once the functionality of individual modules was tested, they were then connected through balancing of molecular quantities such as the ATP:ADP ratio, which should remain globally constant. We then simulated the first minute of a neuron’s life utilising this global model.

During the simulation, the number of cytosolic calcium ions fluctuated periodically displaying cyclic behaviour (Figure 1.C.1.) and the number of calcium ions present in the mitochondria and the ER reached their maximum allowable limit (3.10^7^, Figure 1.C.1. and 1.7.10^9^, Figure 1.C.2, respectively). The number of ATP and ADP molecules, despite oscillations, reached a global dynamic equilibrium state ([ATP]/[ADP] = 1000, [ADP] = 1 mmol.L^-1^, Figure 1.C.3), while the number of ROS increased steadily (Figure 1.C.4); this increase was represented by a step function. The simulation of the iron metabolism yielded similar results to that of the single module simulations (Figure 1.C.5-8), since only a small part of the metabolism was involved in the ROS production, thus linking the system to the rest of the modules. The α-synuclein quantity remained constant with no oligomers or Lewy Bodies being synthesised in this initial stage. This is a particularly encouraging result for the duration of the simulation, since we propose that the increase in α-synuclein molecules would be exclusively due to iron accumulation, which would be triggered by the interaction between ROS and the iron metabolism.

The available information on transition rates allowed us to observe the chain of events leading to protein aggregation, hence triggering the initial disease formation within a minute. This model is sufficiently flexible to allow an adjustment of the rates as more data become available on any of the processes included in the model, particularly on their regulation. We expect such modifications only to extend the timelines to trigger the formation of the disease without disrupting the model structure representing the disease mechanism.

### Case Studies

We investigated two scenarios concerning molecular manifestation of Parkinson’s disease employing this model. The first scenario was based on the assumption that, at some point in the disease, the cytosolic iron exceeds a certain threshold, triggering an abnormal translation of α-synuclein, leading to the formation of detrimental oligomers. The maximal concentration of α-synuclein was observed at 10 µM in SNpc DA neurons (Figure 1.D.1-2). The extent of oligomer and Lewy body aggregate formation simulated by the model was between 10^2^ (oligomers) to 10^4^ (Lewy Bodies) times higher than those suggested in the literature although the model parameters were adopted from the literature (see Table 1 in STAR Methods) (Dong et al., 2018; Iljina et al., 2016). This discrepancy could have been the result of yet unknown or undocumented mechanisms of regulation in the translation or degradation of α-synuclein and its oligomers, which were consequently omitted from the model.

The second scenario investigated the impact of α-synuclein accumulation on ROS production. We tested a mechanism whereby α-synuclein oligomers would affect the antioxidation capacity and reduce the activity of antioxidants. We modeled this inhibition in the model and compared that to the simulation result that omitted any potential interaction of ROS metabolism with α-synuclein. An increase in α-synuclein concentration was observed to trigger an increase in ROS production by an exponential factor in comparison to ROS production, which was not associated with α-synuclein accumulation (Figure 1.D.3.).

## Discussion

We here present a mechanistic model of Parkinson’s disease, which relies on numerical simulation. Although databases on the pathways and interactions pertaining to PD already exist (Fujita et al., 2014), our model is the first ever attempt to incorporate all available literature to construct a mathematical model of the disease.

We showed that a dysfunction in the autonomous pacemaking facilitated by calcium transport may be the main cause for the development of PD in a neuron. Using this model, we were able to demonstrate that the following causality relationships can realistically exist without being irrelevant to or contradictory with available literature, and manifest itself as disease phenotype: High calcium concentration fosters the energy metabolism in the mitochondria. Excessive activation of energy metabolism causes electrons to leak from the mitochondrial transfer chain and generate ROS. Among other responses, ROS is responsible for the dysfunction of iron buffers leading to reactive iron accumulation in the cell. High iron concentration triggers and aggravates α-synuclein translation, leading to its misfolding and accumulation, until the degeneration of the dopaminergic neuron.

The model could be further improved by incorporating the probabilities of developing Parkinson’s disease or of observing randomisation of the SNpc DA neuron pacemaking based on different age groups as extensive epidemiological data becomes available in the future. As more data becomes available on assessing the minimum level of ROS generation to trigger an irreversible disruption of the cell, details on the mechanism relating ROS to α-synuclein could be incorporated into the model to describe the acceleration of neuronal degeneration at late stages of the disease. The model could also be modified to investigate cell-to-cell communication between degenerating and healthy neurons, particularly in relation to ROS production and the prion-like propagation of α-synuclein in the healthy neuron. This could then pave way to studying the microglial implication in the disease by a three-actor model.

SNpc DA neurons have specific characteristics in which they differ from other non-dopaminergic neurons (Chan et al., 2009; Haddad and Nakamura, 2015; Pissadaki and Bolam, 2013; Poewe et al., 2017). Investigating to what extent these specificities account for neuronal degeneration could lead to new discoveries. Moreover, comparing dopaminergic neurons that are not similarly affected during the disease could unveil new mechanisms involved in PD. Indeed, dopaminergic neurons from the ventral tegmental area are far less affected than the ones from the SNpc (Poewe et al., 2017). One known difference is the composition in L-type (Cav1.3) channels at their membrane, which is a promising path to explore (Chan et al., 2007, 2009). As more information becomes available, the model can be modified to represent different neuronal types to allow such investigation.

In conclusion, we propose a flexible, buildable and extendable mechanistic model, which can numerically simulate Parkinson’s disease states. The model is able to handle different initial conditions demonstrating its suitability to be used in personalised medicine, and it not only describes correlations between different events occurring in a neuron during disease onset, but also describes the possible causal relationships between those events. Although causality relationships cannot be proven directly, the model is able to test whether the proposed causalities can exist and if they exist, whether this would be unrealistic or contradictory with regards to available knowledge on disease mechanisms and data in the literature, thus equipping us with a handy tool for investigating yet unexplored hypotheses about Parkinson’s disease, especially those that are challenging to test empirically.

## STAR methods

### Software

The individual models and the connected model are deposited in BioModels (Chelliyah et al., 2015) and assigned the identifier MODEL2001290002. The model was run on Snoopy software (Heiner et al., 2012; Rohr et al., 2010), a dedicated Petri Net simulator handling various Petri Nets extensions written in Java and C++. The proposed complex model involving hundreds of equations and operations during simulation takes 11 minutes to simulate one minute of the whole system’s life utilising standard hardware (CPU (Intel® CoreTM i7, 1.80 GHz) and 16.0 GB RAM).

### Further details on the modelling strategy

The model backbone is described in Results to present the global modelling strategy. The rates of transitions and the concentrations employed in different modules of the model are available in Tables 1 and 2 below. The Petri Net representation of each module is provided as Supplementary Materials. Additional technical details concerning calcium-mediated autonomous pacemaking and iron homeostasis modules, as well as on linking different modules and on the two scenarios investigated are described below.

**STAR Table 2:**
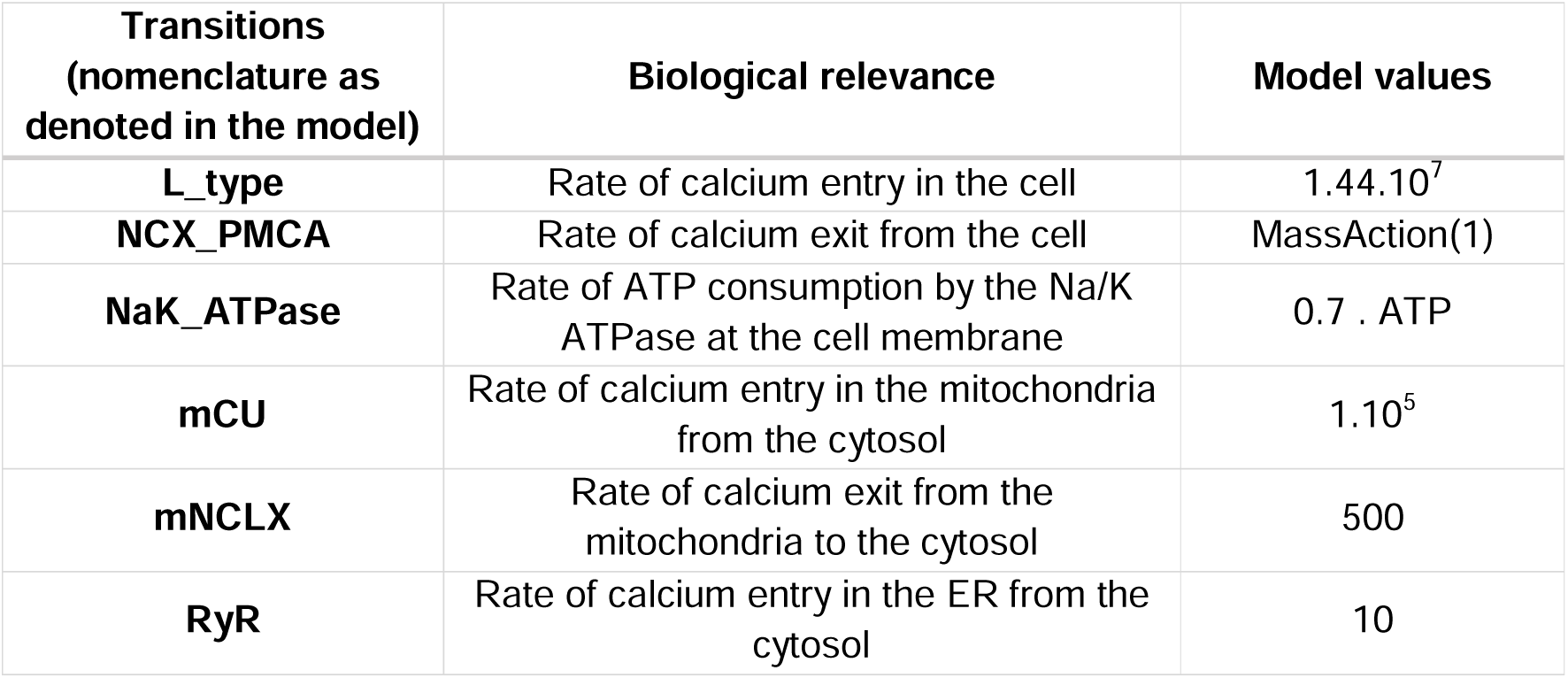
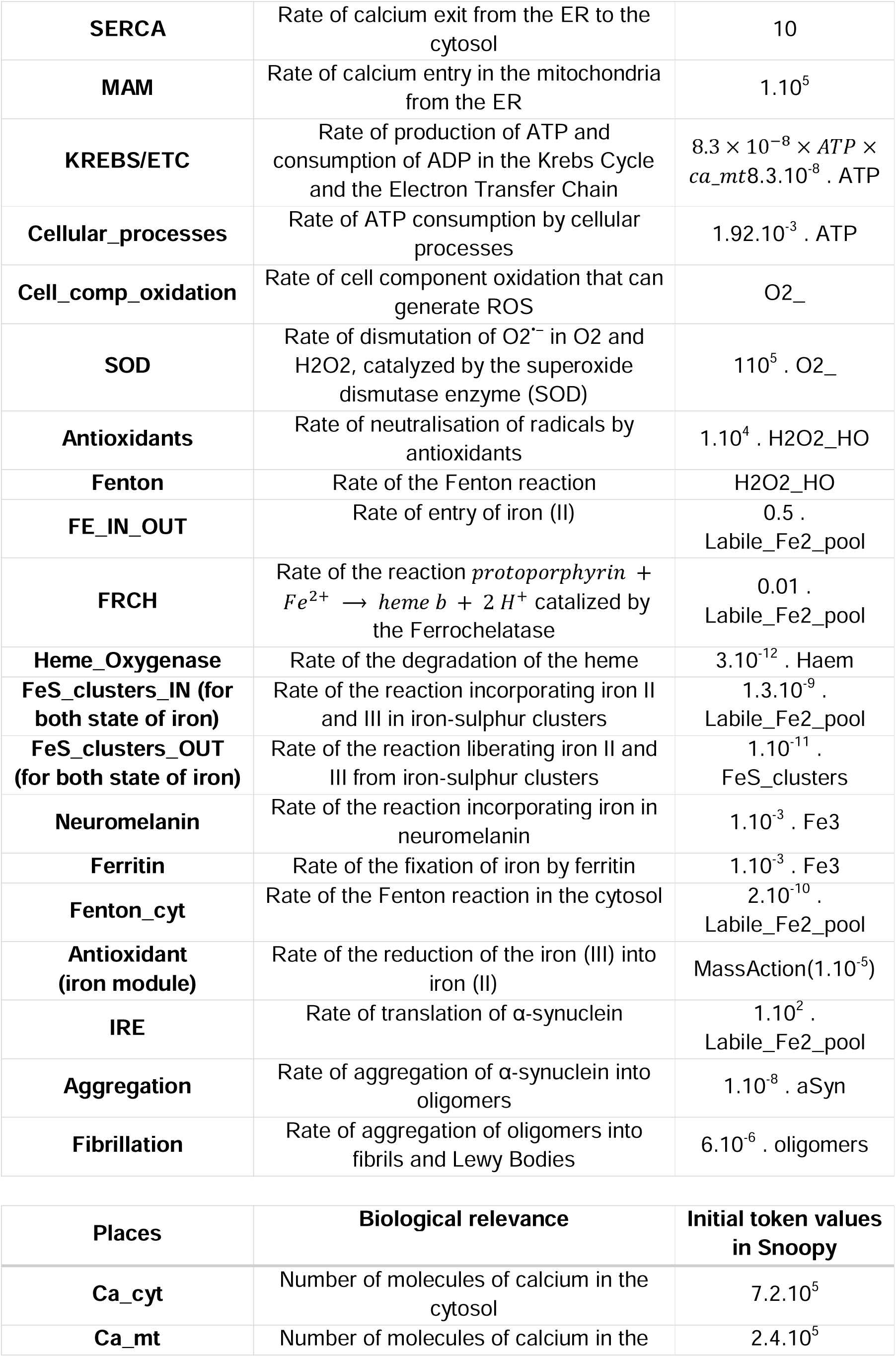
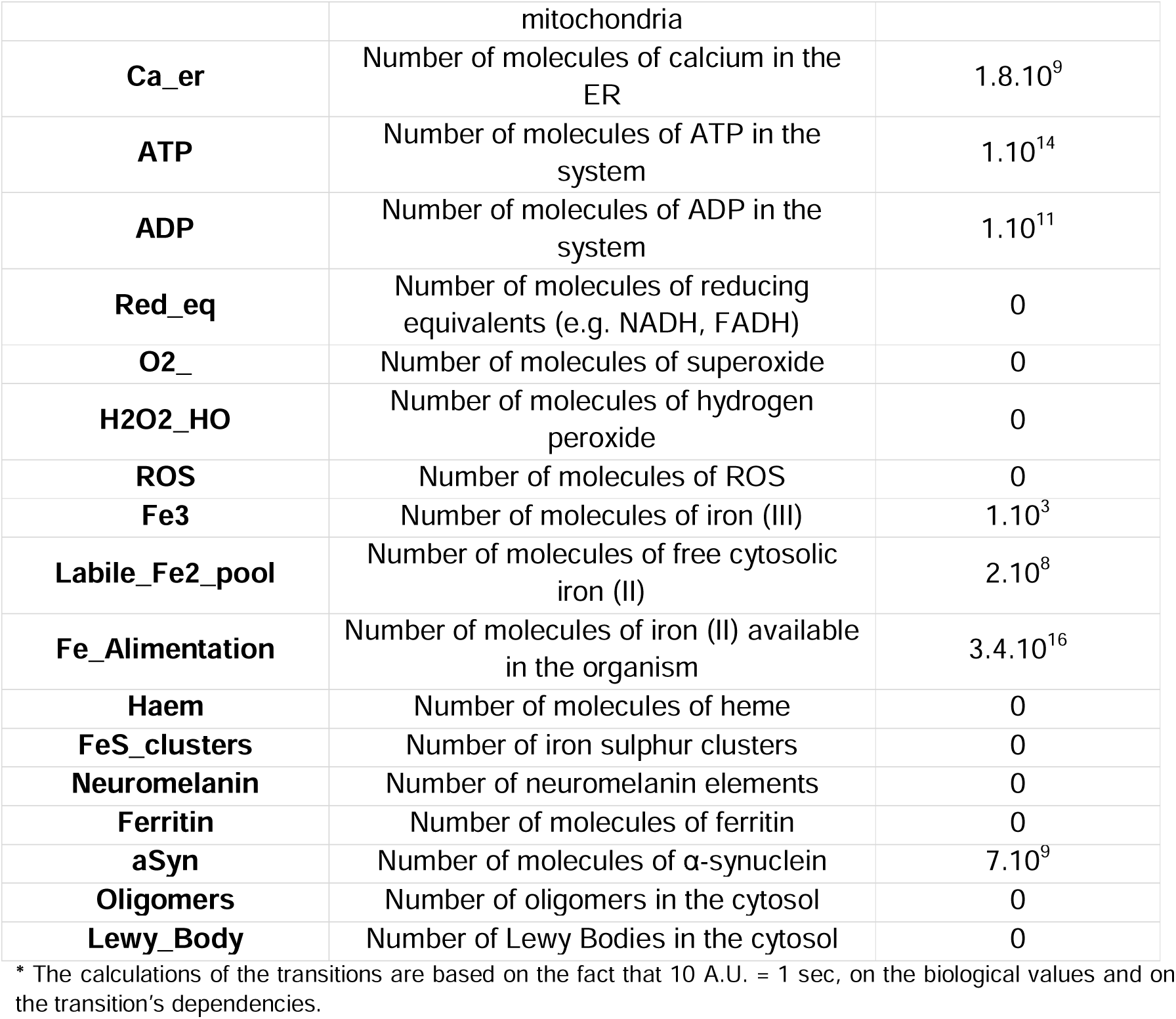
Transitions and initial conditions values used in simulating the complete model.

#### Calcium-mediated autonomous pacemaking

The generation of the autonomous pacemaking is performed by creating a cycle of places, or switches, called on1 to on4, and deterministic transitions that correspond to the delay between each switch, called A to D (Figure S1). When the token is situated in place on1, it remains there for 5 A.U. (transition B), and this event activates the L-type transition, which corresponds to the entry of calcium in the cell. After 5 A.U., the token is moved to place on2 and in this case, there is no time delay (A.U. = 0) (transition C) before it moves to on3. This ensures that there is no delay between the moment the channels that let calcium enter close and the ones that pump calcium out open. Once all the extra calcium has been removed, a latency time of 5 A.U. is allowed (on4 and transition A).

The average time of 1 sec for a spike and the 0.5 sec of latency between each spike were assumed to describe the known feature of natural pacemaking *in silico*. We therefore propose 10 A.U. to represent 1 sec in the simulations. We also engineered the rates such that the known extremal concentrations of cytosolic calcium are not exceeded. All transitions were considered to be deterministic in order to account for the tight regulation around this process. That being said, stochasticity can be introduced in transitions A or C by converting the deterministic transition into a stochastic transition. Stochasticity may be incorporated in the delay between spikes in Transition A, and in the delay between the closing of L-type channels and the opening of calcium pumps in Transition C.

Continuous transitions were employed since rates were available in the literature (STAR Table 1). The stochastic behaviour of the molecules is approximated to be continuous like the concentrations from which they were estimated. Calcium can flow in the mitochondria through VDAC and mCU channels and exit through mNCLX. SERCA is an ER influx pump for calcium, which can exit through the RyR channels. Mitochondrial calcium can be supplied by the ER through the Mitochondrial Associated Membranes (MAM). A threshold concentration of 10 μM is required locally for calcium to enter the mitochondria, and this is represented by read arcs ending on mCU and MAM (Figure S1.a.).

Given the known rates and the maximum concentration of calcium detected into the mitochondria, we introduced an artificial upper bound materialised by the light red inhibitory arcs (Figure S1.a.). Had this not been done, calcium concentration would increase indefinitely, rendering the model biologically irrelevant. The inhibitory arc on RyR, denoted by dark red colour (Figure S1.a.), was created to block the backflow of calcium from the ER during calcium entry. This assumption complies with current literature, but can be revisited in the future as more detailed insight becomes available on calcium homeostasis in the ER.

#### Iron homeostasis

The transitions from the labile iron (II) pool and iron (III) to Fe-S clusters (STAR Table 1, Figure S6) are considered as separate entities despite some Fe-S clusters containing both states of iron, since the tokens in either place can be depleted. Indeed, the nature and definition of Petri Nets would have stopped the transition to fire, upon the depletion of iron (III) for example, disregarding the possibility that Fe–S clusters can also be composed of only iron (II). The green inhibitory arcs are blockers and necessary to represent internal regulation mechanisms. Indeed, the iron cannot accumulate towards infinity into those buffers. The complex biosynthesis and degradation pathways are tightly regulated and involve the intervention of ATP. These blockers are therefore modelling regulatory mechanisms that prevent from overproducing haems, Fe–S clusters, neuromelanin or ferritin by consuming essential resources needed in other cellular processes.

#### Linking the modules

Mitochondrial calcium and the energy metabolism was the first connection to be made. Mitochondria buffer calcium and it is known that calcium ions foster the Krebs cycle by interacting with specific enzymes. Given that the energy metabolism was only accelerated (and not activated) by mitochondrial calcium, we included modifier edges from place named as Ca_mt to KREBS and ETC in order to incorporate the number of mitochondrial calcium molecules into the expression of their rates without consuming them. KREBS and ETC have exactly the same rate in order to keep the overall number of molecules of ATP and ADP constant, however, the molecules of ATP and ADP are also involved in SERCA pumping and at the membrane for establishing ions gradients. Therefore, the transition representing sodium potassium ATPase pump (NaK_ATPase, Figure 1.B.) at the membrane was modified from an instantaneous transition to a continuous transition where the number of molecules of ATP and ADP were also considered as continuous. The transition is activated at the end of the cytosolic calcium spike by a read arc. This read arc allows for the simulation of a continuous rate of ATP consumption and ADP production. The transition is consuming 70% of the ATP at time *t*, instead of 70% of the ATP at time *t* = 0, in order to keep the conservation of the number of ATP and ADP molecules.

The ETC transition was connected to the synthesis of O2°^-^ (O2_, Figure 1.B.) to account for the leaking of electrons from the electron transfer chain causing the formation of radicals by a reaction with dioxygen. Limited data are found on the frequency of the ETC leak; therefore, it was approximated as one electron per cycle, only to be revaluated if ROS accumulation would be too fast or as new data becomes available in the future. The labile iron pool was associated with the ROS module through the Fenton Reaction. We consider that all ROS produced during the Fenton Reactions and by the iron homeostasis module are modelled in the ROS module and is represented by a single rate of transition. A final link was established between the iron metabolism and the α-synuclein protein. Since the concentration of the protein was maintained constant prior to activation by iron, the rate of its translation (denoted as IRE) would thus be activated as a given concentration/number of molecules of iron (II) become available in the place denoted as Labile_Fe2_pool. This number was fixed at an intermediary value, which was arbitrarily chosen and was between the iron concentration in healthy neurons and those found in PD brains, closer to that of former than the latter.

### Case study 1

In this scenario the activation of α-synuclein translation upon iron accumulation reaching a critical value followed by protein aggregation into oligomers and fibrils that can form Lewy Bodies was investigated. The average concentration of oligomers found in PD brains was reported to be at around 10 pM (Hansson et al., 2014; Horrocks et al., 2016; Hughes et al., 2019). This concentration yields approximately 100 to 1000 oligomers corresponding to 1 to 10 Lewy Body/fibrils in our model. We consider the iron concentration in the pool to remain constant. The rate of translation was set 10 times that of the estimated biological rate in order to simulate a longer period of time. Indeed, instead of ∼ 2.7h, we will be able to study the evolution of α-synuclein, oligomers and Lewy Bodies during about a day (Figure 1.D.1-2). The maximum number of α-synuclein proteins was set as 5.10^10^ molecules based on available literature.

### Case study 2

This scenario was modelled by linking the oligomers to the antioxidants (ROS module) via modifier edges and running the simulation over the complete model. The number of oligomers was taken into account into the rate of neutralization of radicals by antioxidants and the values were normalized to remain within acceptable limits of the global initial rate of ROS accumulation.

## Acknowledgements

The authors would like to thank Maria Zacharopoulou for useful discussions on disease pathophysiology.DD gratefully acknowledges the funding from the Leverhulme Trust and the Isaac Newton Trust (ECF-2016-681). MN was partially supported by an Erasmus+ grant, allotted by the Ecole Normale Supérieure de Paris – PSL University, 75005 Paris, France. G.S.K.S. acknowledges funding from the Wellcome Trust, the UK Medical Research Council (MRC), Alzheimer Research UK (ARUK), and Infinitus China Ltd.

