## Supplementary Material for "Formal model of Parkinson’s disease neurons unveils possible causality links in the pathophysiology of the disease"

**Supplementary materials**

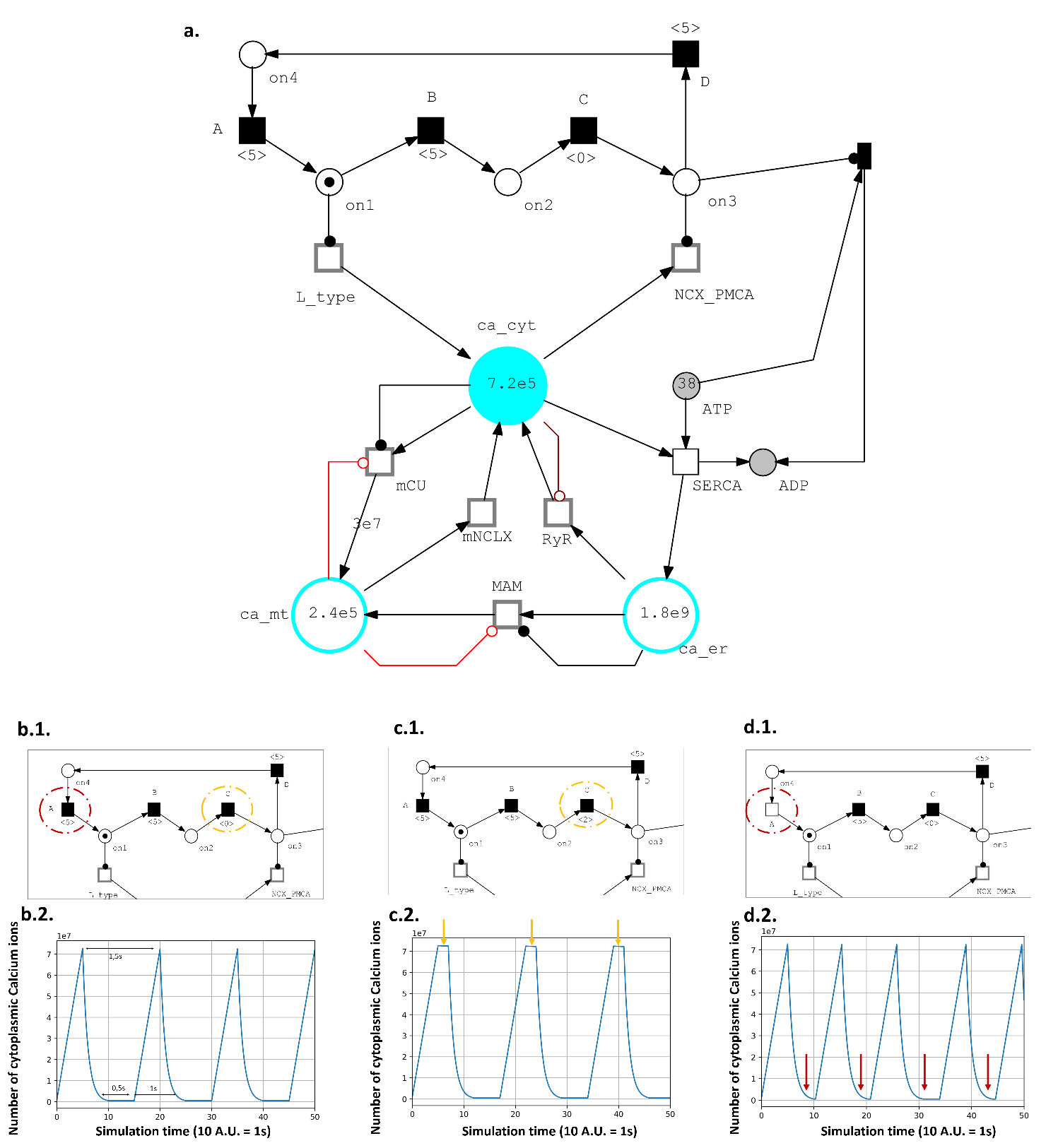
**Calcium metabolism module**

**Figure S1: Calcium Petri Net Module and its adaptation to ageing individuals. a.1**. represents the model used in the project. It consists in the pacemaking generation through a system of switches (places, on1 to on4, with one possible token and deterministic transitions, A to D). 10 A.U. of simulation time correspond to 1s. The cytoplasmic calcium can then be loaded in the mitochondria or the ER. **a.2.** The generated pacemaking, expressed in number of calcium ions, corresponding to the biological system**. b.** A change in the delay of transition C affects the behaviour of the system by longer exposing the cell to a high concentration of calcium. **c.** Converting the transition A into a stochastic one permits to mimic the behaviour of aged neurons and add randomness into the delay between two spikes.

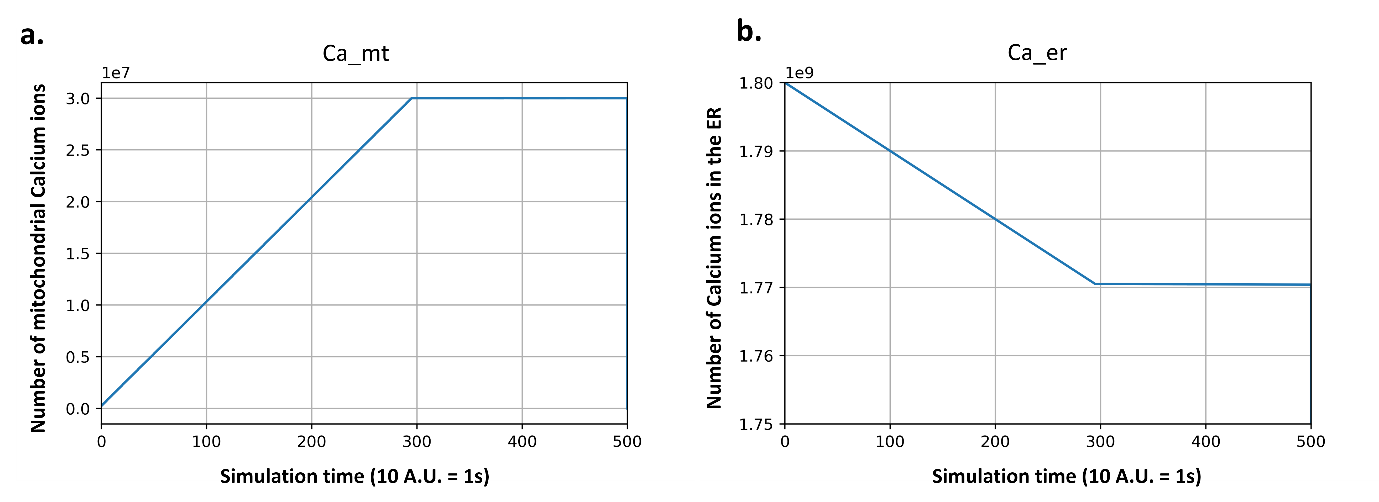

**Figure S2: Calcium concentration evolution in the first 50 seconds of simulation. a. Mitochondrial calcium.** The transient state corresponds to an increase of calcium in the first 30 sec until it reaches its higher concentration and stays in its steady state**. b. ER calcium.** The concentration of calcium decreases quickly until the mitochondria has reached its maximum threshold. Then the concentration decreases at a very low rate and finally reaches a steady state.

**Energy metabolism module**

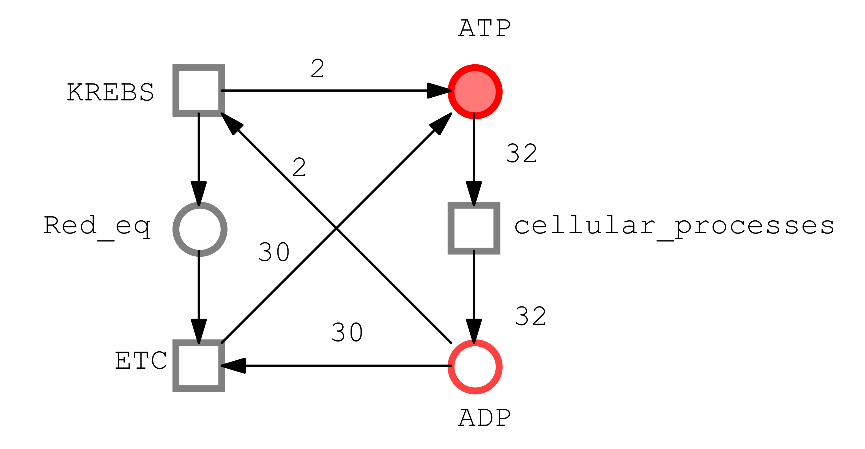

**Figure S3: Energy Metabolism Petri Net Module.** We consider that because of the electron leak, only 32 ATP are finally synthesised in the cell. The Krebs cycle contributes to 2 ATP and the reducing equivalent necessary to the ETC to synthesise the last 30 ATP. The ATP is then consumed by cellular processes. Extra care had to be given to the rates because the number of molecules of ATP and ADP has to be conserved.

**ROS module**

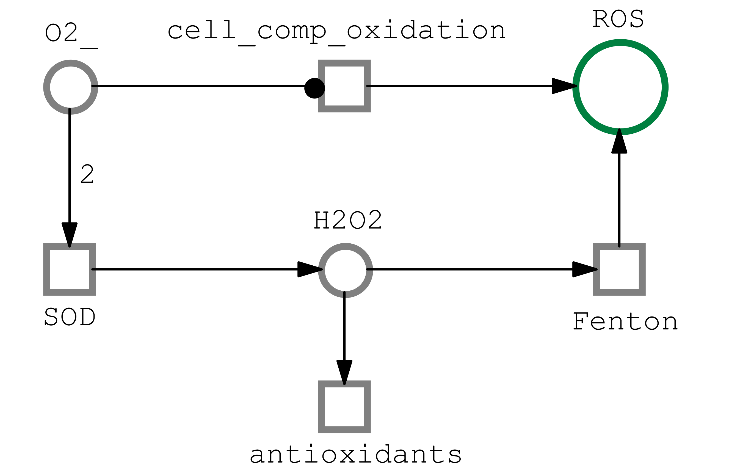

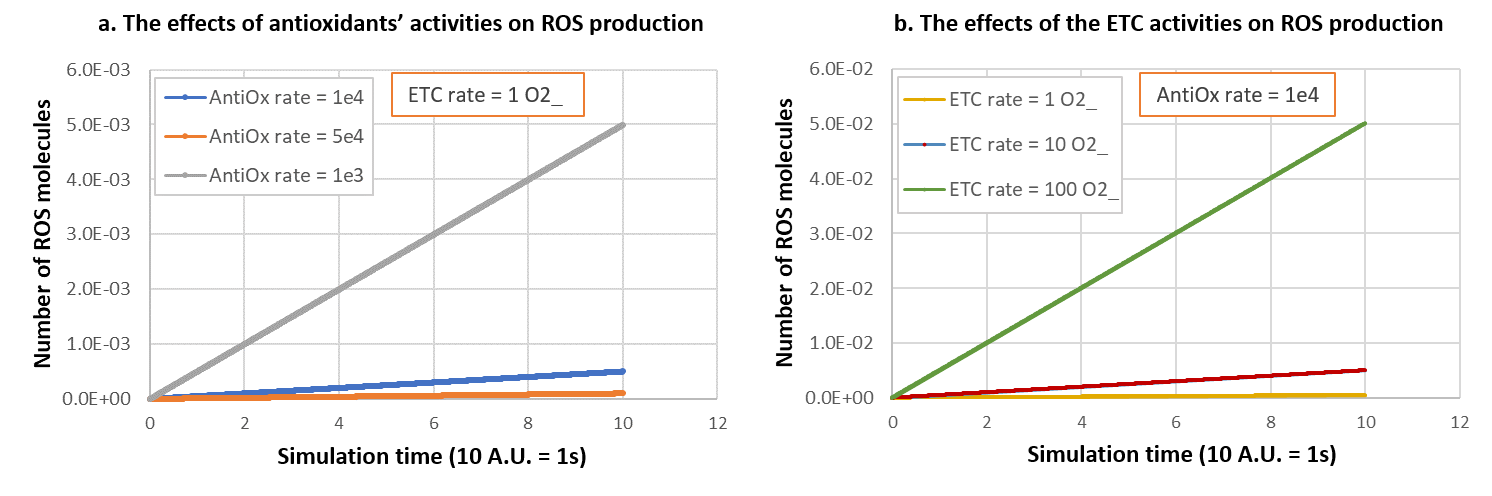
**Figure S4: ROS Petri Net Module.** O2_ (O2°^-^) is generated by the ETC and, at a certain concentration, can oxidise cell components, it is thus considered as a source of ROS. It can also be reduced into H_2_O_2_ (H_2_O_2_ and O_2_) and the hydrogen peroxide is generally safely discarded by antioxidants, although a little fraction of it can produce ROS.

**Figure S5: Effects of the antioxidants’ and the ETC rates on the production of ROS in Module 3. a. The effects of antioxidants’ activities on ROS production.** At a fixed ETC rate of 1 O2_ molecule per A.U., the number of ROS and their production speed increases dramatically with the augmentation of the antioxidants’ rate (AntiOx rate). **b. The effects of ETC activities on ROS production.** At a fixed antioxidants’ rate of 1.10^4^ discarded molecules per A.U., the number of ROS and their production speed increases with the augmentation of the ETC rate, with a similar tendency compared to **a.**

**Iron metabolism module**

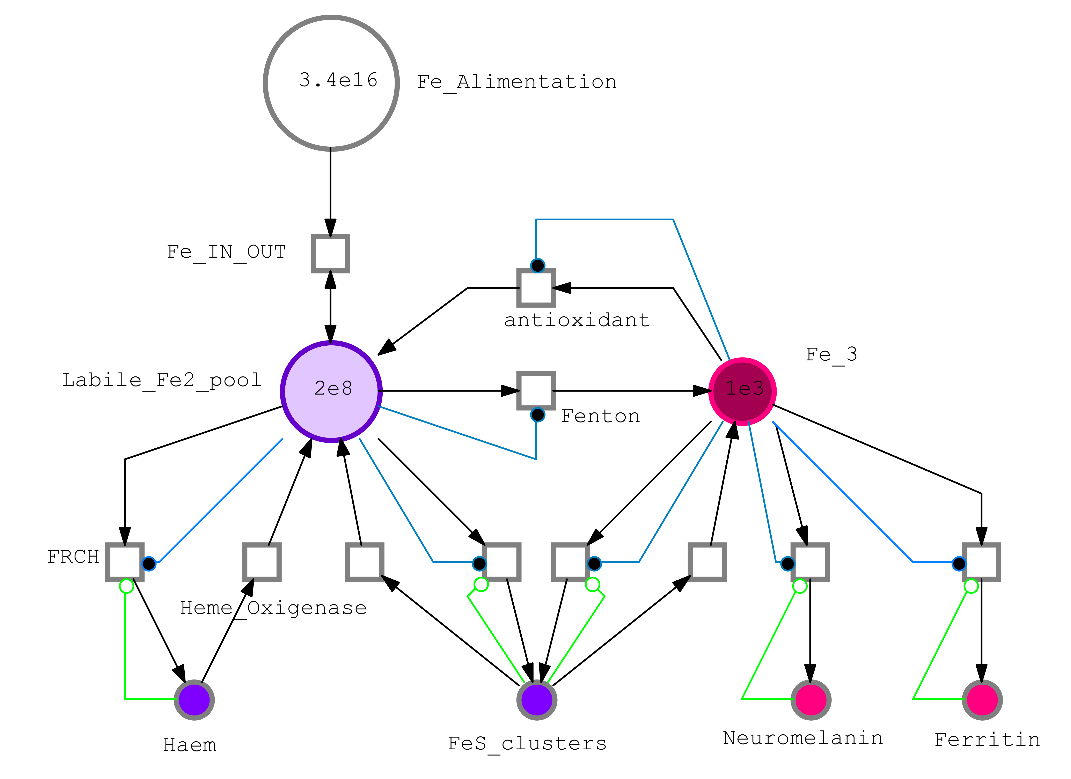

**Figure S6: Iron metabolism Petri Net Module.** The iron is supplied by the daily intake Fe_Alimentation under the state of iron (II), Labile_Fe2_pool. It can be directly available for reactions or be implicated in Haem and FeS_clusters synthesis. Iron (II) can either be oxidised to iron (III) which is highly regulated because of its propensity to react with cellular components, and stored into FeS_clusters, Neuromelanin or Ferritin.

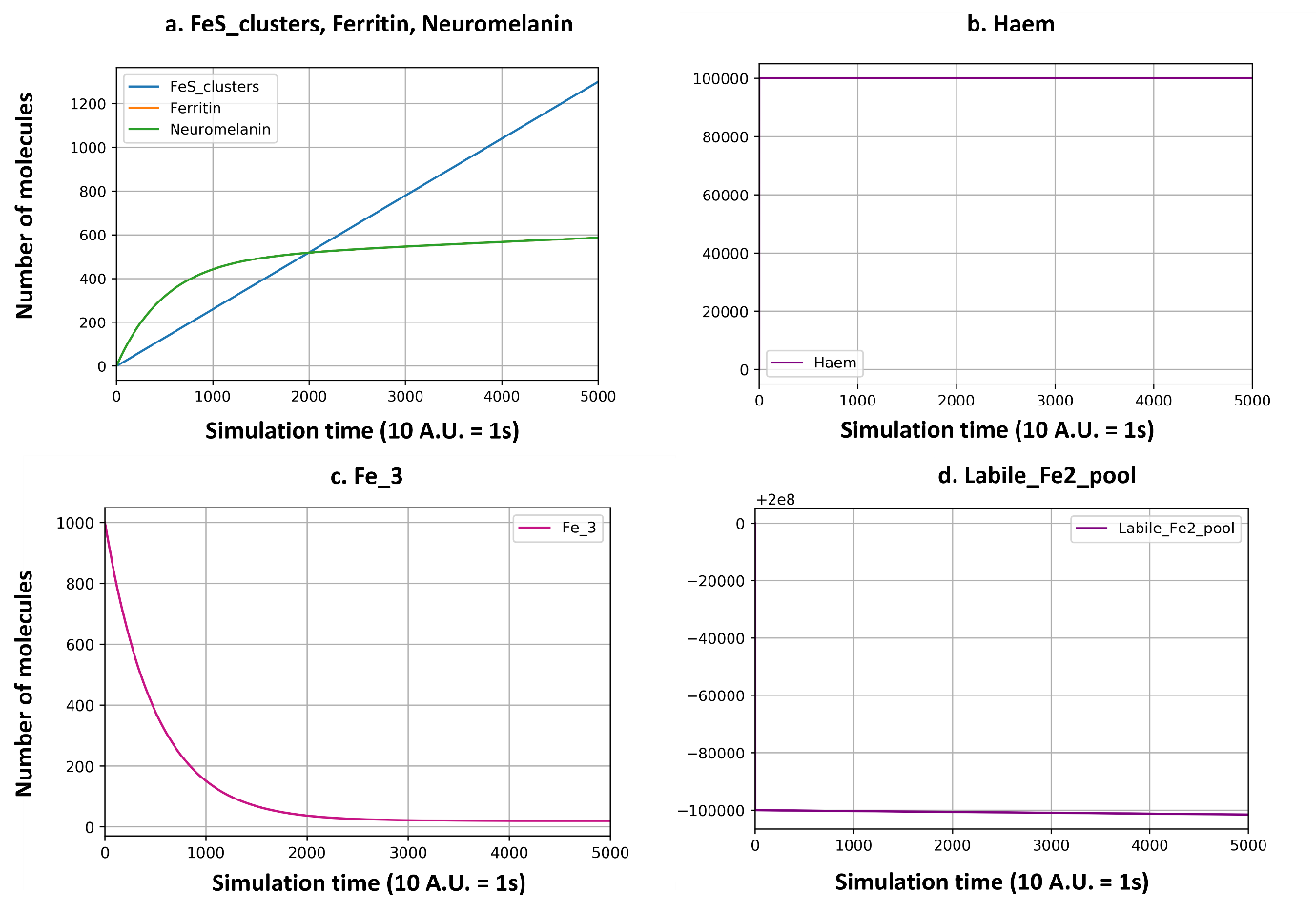
**Figure S7: Evolution of the quantities of iron species during the first 500 sec of simulation. a. Fe–S clusters, Ferritin and Neuromelanin evolution.** Ferritin and Neuromelanin have the same rate of iron (III) incorporation. Because of the internal regulation, they are getting close to a certain threshold (~600 molecules). FeS_clusters are still increasing, because of a higher quantity threshold. **b. Haem evolution.** The rate of haem synthesis is high. All the haems necessary are synthesised in the first instants of the simulation. **c. Fe^3+^ evolution.** The iron (III) reaches a low number of molecules steady state. **d. Labile iron pool evolution**. Once all the iron is used for the essential cell synthesis, it decreases very slowly and is expected to reach a steady state. The diminution is represented by the increment of negative numbers to 2.10^8^.

**α-synuclein module**

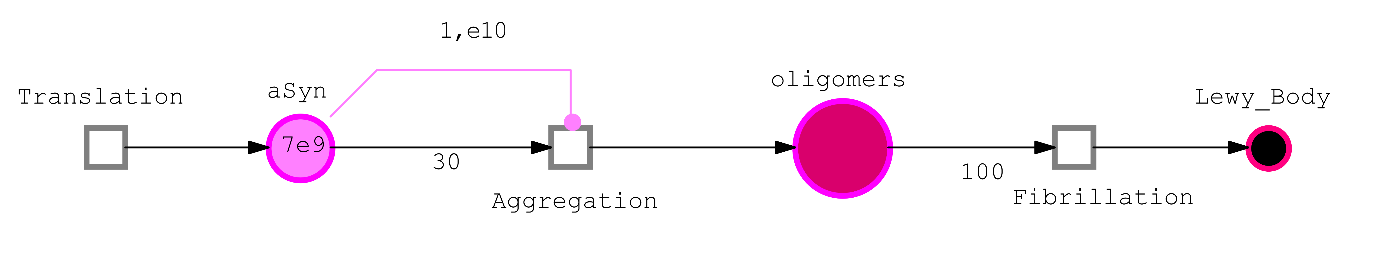

**Figure S8: α-synuclein Petri Net module.** α-synuclein (aSyn) is translated and can aggregate if its number of molecules exceed 1.10^10^. 30 molecules are then used to form one oligomer which can in turn be aggregated into fibrils or into a Lewy_Body.
